# On the evolution of neural decisions from uncertain visual input to uncertain actions

**DOI:** 10.1101/803049

**Authors:** Alessandro Tomassini, Darren Price, Jiaxiang Zhang, James B Rowe

## Abstract

Behavior can be conceived as the result of a sequence in which the outcomes of perceptual decisions inform decisions on which action to take. However, the relationship between these processes, and spatiotemporal dynamics of the visual-to-motor transformation remains unclear. Here, we combined accumulation-to-threshold models and electro-magnetoencephalography, to trace neural correlates of sensorimotor decisions in space, time and frequency. We challenge the assumption of sequential decisions, with evidence that visuomotor processing unfolds through a continuous flow of information from sensory to motor regions. Action selection is initiated before regional visual decisions are completed. By linking behavior and physiology through theoretical decision models, we identify simultaneous forward and backward flow of information for visuomotor decisions between sensory and motor regions, in beta and gamma ranges. The model of integrated visuomotor decisions provides a powerful approach to investigate behavioral disorders that impair the ability to use sensory inputs to guide appropriate actions.

## Introduction

Human behaviors are the result of many decisions, from early or automatic perceptual inferences about our environment to complex goal-directed choices between alternate courses of action. Three broad lines of research have made separate contributions to understanding such decisions. First, the psychophysical analysis of visuomotor task performance and reaction times, in health^1^ or in the presence of focal^2^ and degenerative brain lesions^3^. Second, the functional anatomical analysis of decision making using brain imaging and neurophysiology, including paradigms that manipulate visual uncertainty^4^, action selection^5^, or outcome evaluation^6^. Third, the development of computational models of how decisions can be reached, at the level of neuronal ensembles^7^ or groups of individuals^8^.

It remains a challenge however, to bring these separate lines of enquiry together in a unified model of neurophysiologically informed decision process, embedded in a functional anatomical framework, to explain the transformation of noisy visual inputs to alternative motor outputs. The anatomical framework has an additional requirement, which is to accommodate the evidence for functional segregation between sensory and motor areas at the same time as allowing the flow of information through hierarchical and distributed brain networks. Here we propose an integrated account of visuomotor decision-making, as summarised in Fig. 1, working from a novel visuomotor task that adjusts sensory and action uncertainty during functional brain imaging by combined electro-/magnetoencephalography (MEEG).

**Figure 1.**
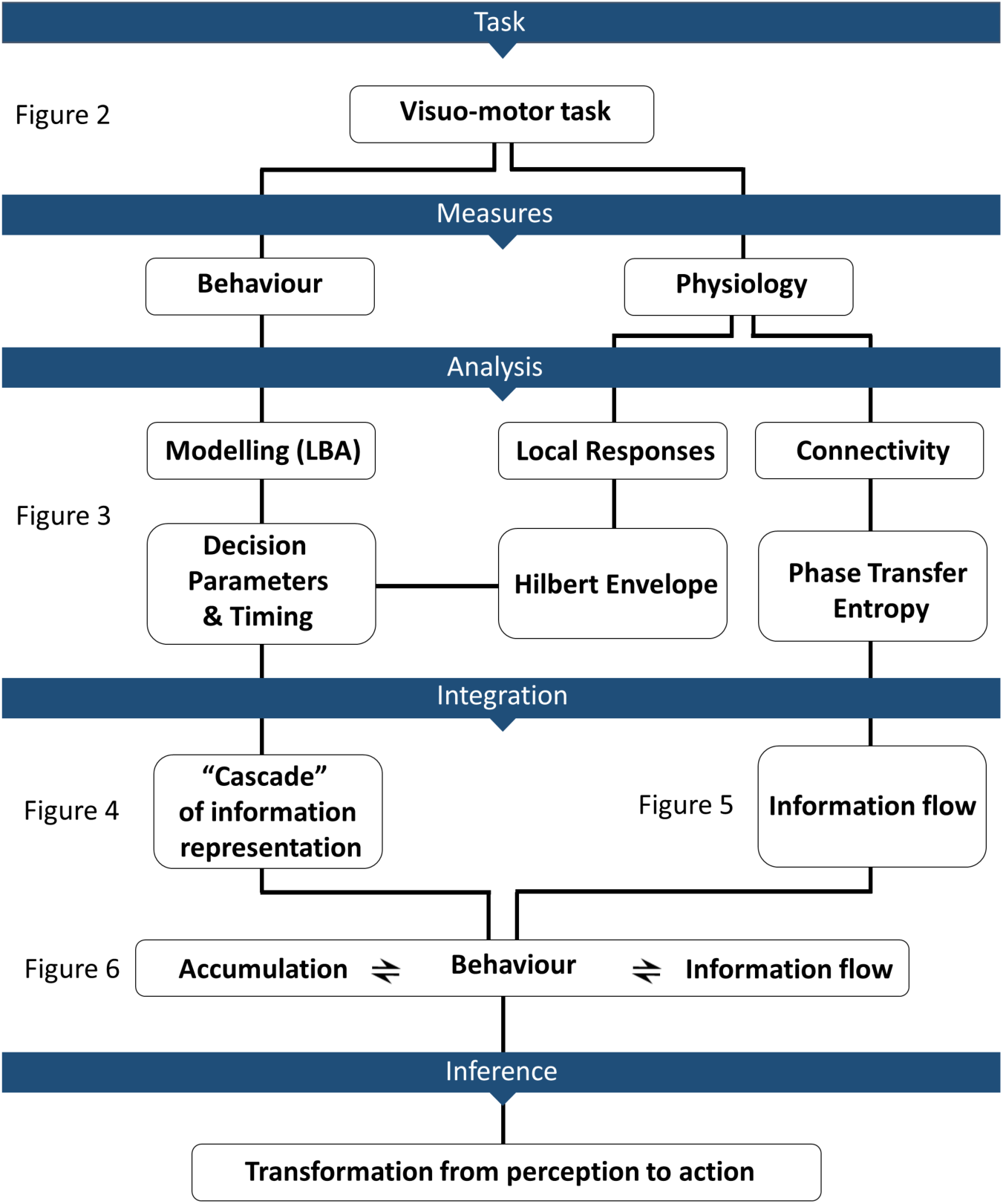
Overview of the study. We combined behavioural, computational, and neuroimaging approaches to provide an integrated perspective of the decision processes linking perception to action. Each section is expanded in a subsequent figure, as directed.

In mathematical psychology it has been argued that decisions and their latencies are controlled by when the cumulative evidence in favour of a choice reaches a criterion decision threshold^9^. We identify the accumulation-to-threshold of latent variables representing sensory evidences, based on the transformation of visual signals into evidence about the behaviorally relevant stimulus features (perceptual decisions^10,11^); and the analogous *evidence* for motor schema, which have been termed motor intentions (action decisions^12,13^).

Rather than arbitrarily divide visual from motor transformations, we investigated their associated decision processes by separately manipulating uncertainty in the identity of visual features (perceptual uncertainty^11^, by variable motion coherence) and range of possible actions (action uncertainty^13,14^, by variable number of response options).The task combined elements of the classic motion discrimination task^11^ with a response selection task^13^: noisy visual stimuli indicated the one or more response options, which were executed by pressing a corresponding button (**Fig. 2**). By manipulating motion coherence in the random dots stimuli and the number of offered choices, we sought to isolate the neural signatures of decision-evidence accumulation for perceptual and action decisions, respectively.

**Figure 2.**
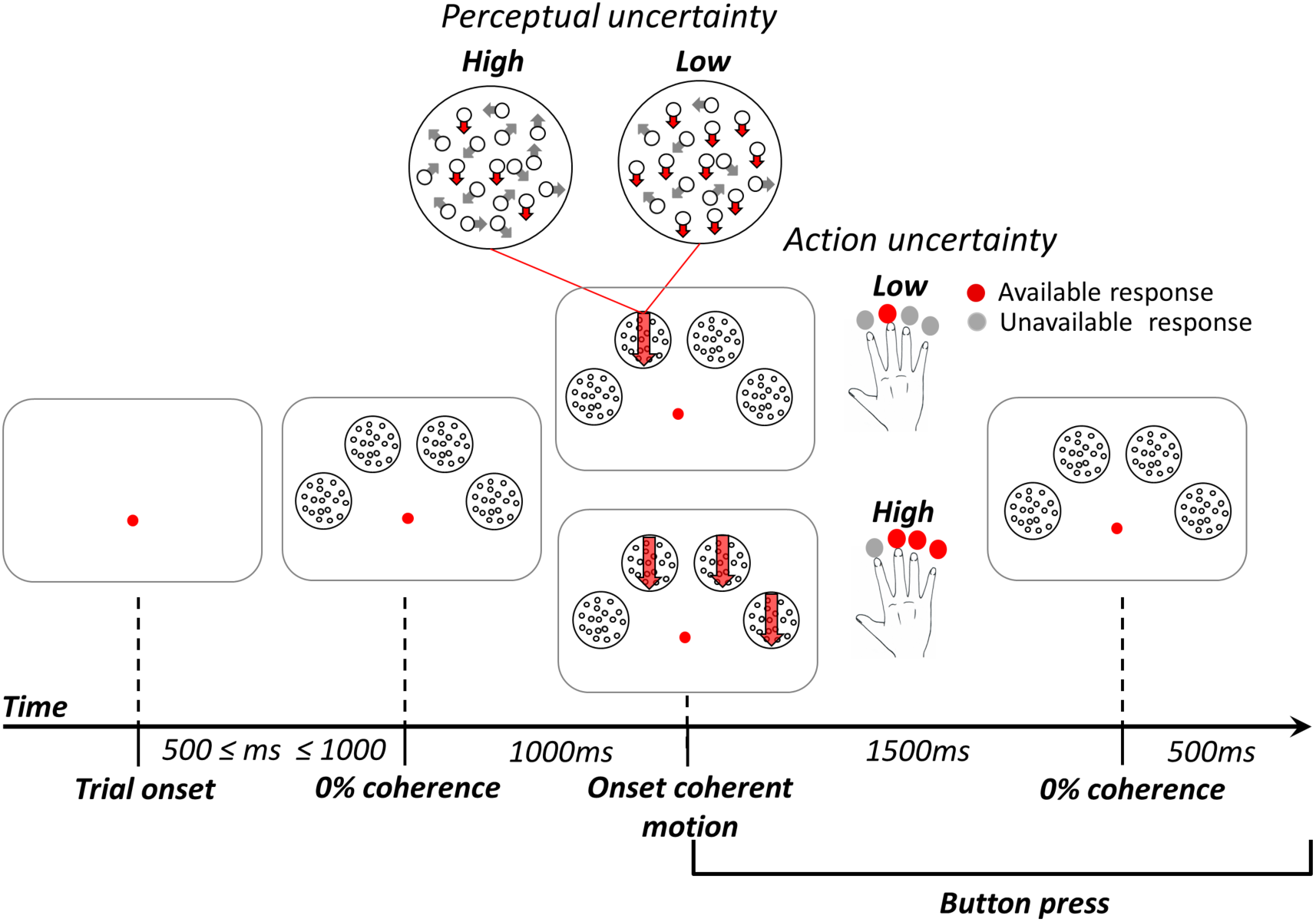
Experimental manipulation of perceptual and action uncertainty. Participants pressed the button corresponding to the coherent stimulus (red downward arrow). When there were more than one coherent stimulus, they selected one response and pressed the corresponding button. Perceptual uncertainty was manipulated by changing the coherence of dot motion (i.e. by changing the motion strength), whereas action uncertainty was manipulated by changing the number of available options (i.e. the number of coherent stimuli to choose from). Perceptual and action uncertainty varied across trials in a 2 by 2 factorial design.

Several brain regions have been identified that accumulate perceptual evidence^10,11^ and motor intentions^12,13^. However, it is necessary to understand how a network of accumulator regions orchestrates their activity for the critical transformation between perceptual and action decisions. Specifically, we sought to distinguish (i) a serial process^15^ where perceptual decisions are complete and their output passed to motor accumulators, from (ii) a continuous flow of information^16^, through perceptual to associative and motor regions before completion of perceptual analysis. A serial process would be robust to error, but continuous flow would enable faster action decisions. To differentiate these alternatives, we mapped the modelled temporal profile of evidence accumulation to MEEG induced power^17^, trial-by-trial. The temporal evolution of predicted evidence was based on behaviorally optimized generative linear ballistic accumulator model of the decision.

We focused on the beta and gamma band power as the candidate correlates of the evidence for three reasons. First, the growing evidence for separate functions of gamma and beta in the feedforward and feedback of information respectively in hierarchical brain networks^18^. Second, that the accumulation of evidence for perceptual choices correlates with gamma-frequency oscillations^17^. Third, that the processes underlying the deliberation between alternate actions have been associated with beta power modulation^19,20^.

The use of MEEG affords a source model of cortical generators^21^ and enables the functional segregation of sensory and motor area, as well as areas where sensory-motor transformations occur. Complementary connectivity measures (phase transfer entropy^22^) reveal the flow of information between areas, orchestrating the emergence of decision-evidences across decision networks.

We show that evidence accumulation in motor and prefrontal cortex begins very soon after visual cortex, and before perceptual decisions are concluded. We further demonstrate that the timing of evidence accumulation and the direction of flow of information between widespread sensory, motor and association cortices differ between Beta (13-30Hz) and Gamma (31-90Hz) frequency range. An early sweep of Gamma activity across an occipito-parietal-frontal network precedes the gradual arising of Beta mediated decision signals.

These signals emerge progressively in a lateralized caudo-rostral cascade unfolding along the dorsal stream. The cascade is mainly driven by a continuous flow of information from posterior visual areas to anterior action control regions. Crucially, the strength of the information flow (as measured by phase-transfer entropy) determines the speed of progression throughout the stages of information processing from perception through action as reflected by a positive relationship between connectivity and both faster model accumulation-rates and shorter reaction-times. This provides an important formal link between behaviour, established models of decision-making, and connectivity measures. Taken together, the results reveal a continuous flow of information transmitted and integrated through a hierarchical network that transforms decision-making from perception to action.

## Results

### Uncertainty modulates the speed of decisions

Participants performed the task first in a training session where individual motion thresholds were estimated for both low and high action uncertainty levels (**Fig. 3a**). Subsequently, participants performed the task with the motion thresholds that standardized performance, while undergoing MEEG scan.

**Figure 3.**
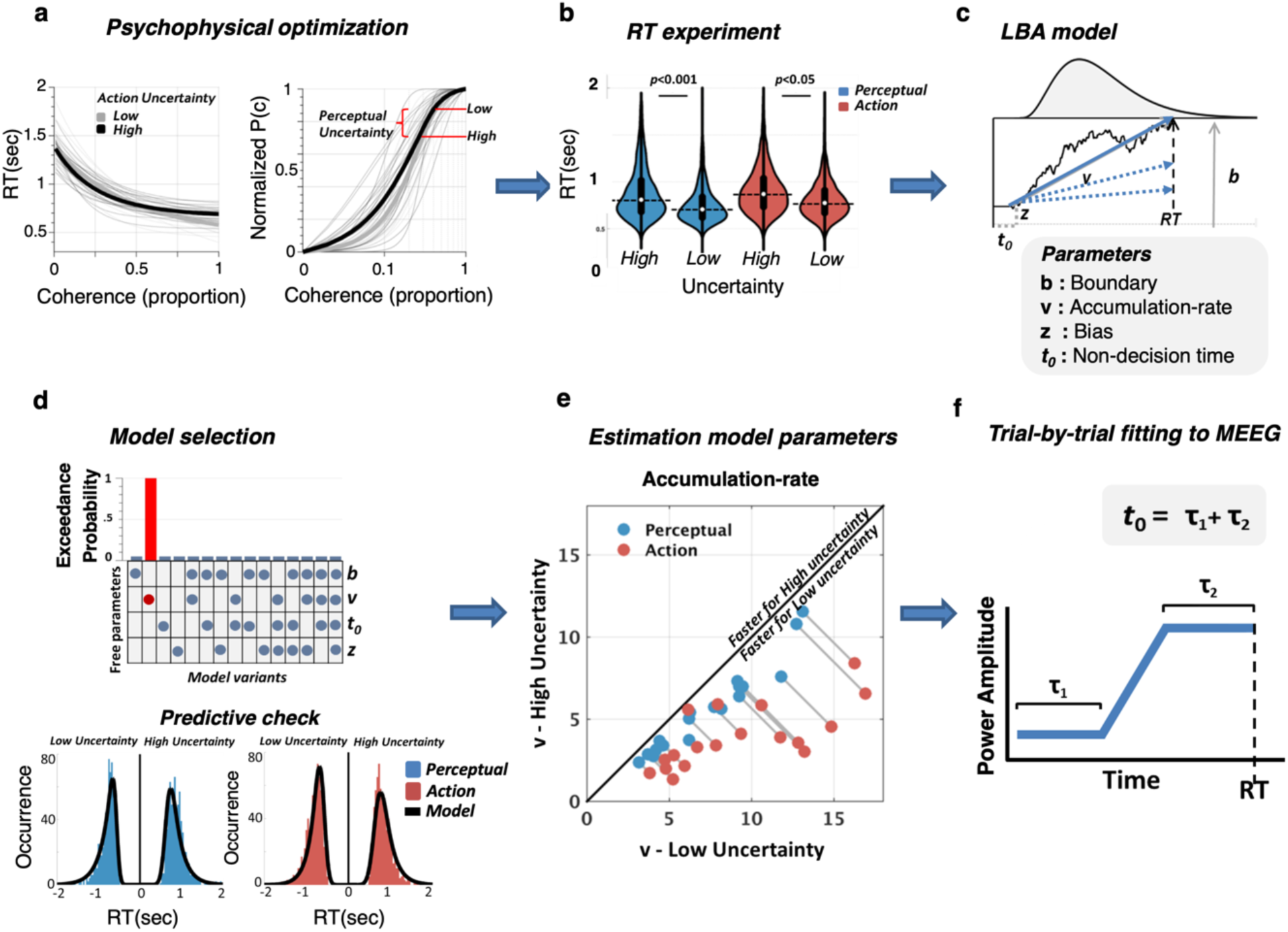
Uncertainty modulates reaction times and the speed of decision evidence accumulation. **(a)** Decision evidence titrated to the participants’ individual motion sensitivity. Reaction times (left panel) and accuracy (right panel) varied with motion strength (grey lines: individual data, black tick lines: mean data). Low and High perceptual uncertainty were estimated at the 75% and 90% accuracy levels, respectively. **(b)** During the experiment, reaction times were significantly modulated by both perceptual and action uncertainty (aggregated data for uncertainty type, rm-ANOVA on log(RT)). **(c)** The task was modelled using a race accumulation model (Linear ballistic accumulator, LBA). Noisy evidence accumulates over time at a rate *v* up to a decision bound *b*. The fastest accumulator (thick blue arrow) determines the choice. Non-decision time linked to sensorial and motor processes (*t0*) sums to the evidence accumulation time to produce reaction-times. **(d)** Bayesian model comparison: changes in the sole drift-rate best accounted for the behavioral data. The quality of fit is also seen by the overlap of empirical reaction time distributions for each condition, with data simulated using the winning model. **(e)** The model predicted faster accumulation of decision evidence when uncertainty is low for both action and perceptual uncertainty (grey lines connects data points from each participant). **(f)** Model predicted activity was fitted to the power envelope of the MEEG signal in a trial-by-trial fashion to identify the latencies of accumulators of decision evidence. Non decision time (*t0*) was decomposed into pre-accumulation (*tau1*) and post-accumulation (*tau2*) time reflecting perceptual and motor processes, respectively.

Behavioral performance scaled with levels of perceptual and action uncertainties, confirming the desired effect of task manipulations. During training, participants where slower and less accurate when motion coherence was lower (**Fig. 3a**). Similarly, during the scan session (**Fig. 3b** and **SI Appendix, Fig. S1**) responses were slower under high perceptual (low = 0.77s ±0.1; high = 0.88s±0.1; F(1,17) = 158.17 p < 0.0001; post-hoc p < 0.0001; ηp^2^ = 0.902) and action uncertainty (low = 0.80s ± 0.13; high = 0.85s±0.1; F(1,17) = 6.28 p = 0.022; post-hoc p = 0.022; ηp^2^ =0.269; 2-by-2 repeated measures ANOVA; Tukey-Kramer correction).

Participants’ choices were largely independent over trials as indicated by Shannon’s equitability index (mean 0.77 ± 0.016 SD, see SI Appendix and Fig. S2 for statistics) calculated for sequential choice pairs^13^.

### Uncertainty modulates the rate of evidence accumulation

To decompose the behavioral performance into cognitively relevant latent variables, we fitted accumulation-to-threshold models (Linear Ballistic Accumulator^23^, LBA), to each participant’s reaction time and accuracy data.

Uncertainty can slow responses by reducing the speed of information accumulation (accumulation-rate), increasing response caution (decision boundary), stretching the time required by perceptual and motor processes not directly related to the decision process (non-decision time), or by a combination thereof. To differentiate these competing mechanisms, we fitted all possible combinations of free parameters in a set of 15 LBAs. We compared the goodness-of-fit of each model using random-effects Bayesian model comparison^24,25^. The model comparison revealed that changes in the accumulation-rate alone (model number 2; **Fig. 3d** top panel) accounted parsimoniously for the effects of uncertainty on behavior. The goodness-of-fit of the winning model was further confirmed by posterior predictive checks (**Fig. 3d** bottom panel), performed by simulating data under the winning model and then comparing these to the observed data.

In the winning model (henceforth, the LBA model), high uncertainty is associated with comparatively slow accumulation rates. This relationship between uncertainty and accumulation rate held for both perception (z = 3.723, p = 0.00019; Wilkoxon sign rank test) and action (z = 3.723, p = 0.00019; Wilkoxon sign rank test) uncertainty, as well as for each subject (**Fig. 3e**), in accord with previous studies^11,14^.

Non-decision time (*t0*), encompassing sensory delays and motor execution, was estimated to be 370ms on average (see **SI Appendix, table S1**), which is within the plausible range of non-decision times for humans^26^.

### Localization of decision-evidence accumulation

The temporal evolution of the combined MEG and EEG spectral power (power envelope) from each region of interest (ROI) of the parcellated cortical surface served as the signal for our analysis in beta (13-30Hz) and gamma (31-90Hz) bands. The time onset of evidence accumulation across ROIs was identified by optimizing the split of the non-decision time before and after the accumulation period using Spearman correlation to the single-trial MEEG power envelope (**Fig. 3f**, see **Methods and Materials for details, supplemental Fig. S3** for the statistical map). This allows one to depict in space and time the emergence of decision-evidence accumulation.

In support of our hypothesis that both increasing and non-increasing activity can mediate evidence accumulation^27,28^, we found significant (negative) correlations between the LBA model predictions and the MEEG oscillations in beta and gamma bands^17^ (**Fig. 4**). Specifically, for both beta and gamma, neural activity after coherence onset desynchronized in a graded fashion and peaked approximately before response suggesting a form of threshold mechanism (**Fig. 4a**)^29,30^.

**Figure 4.**
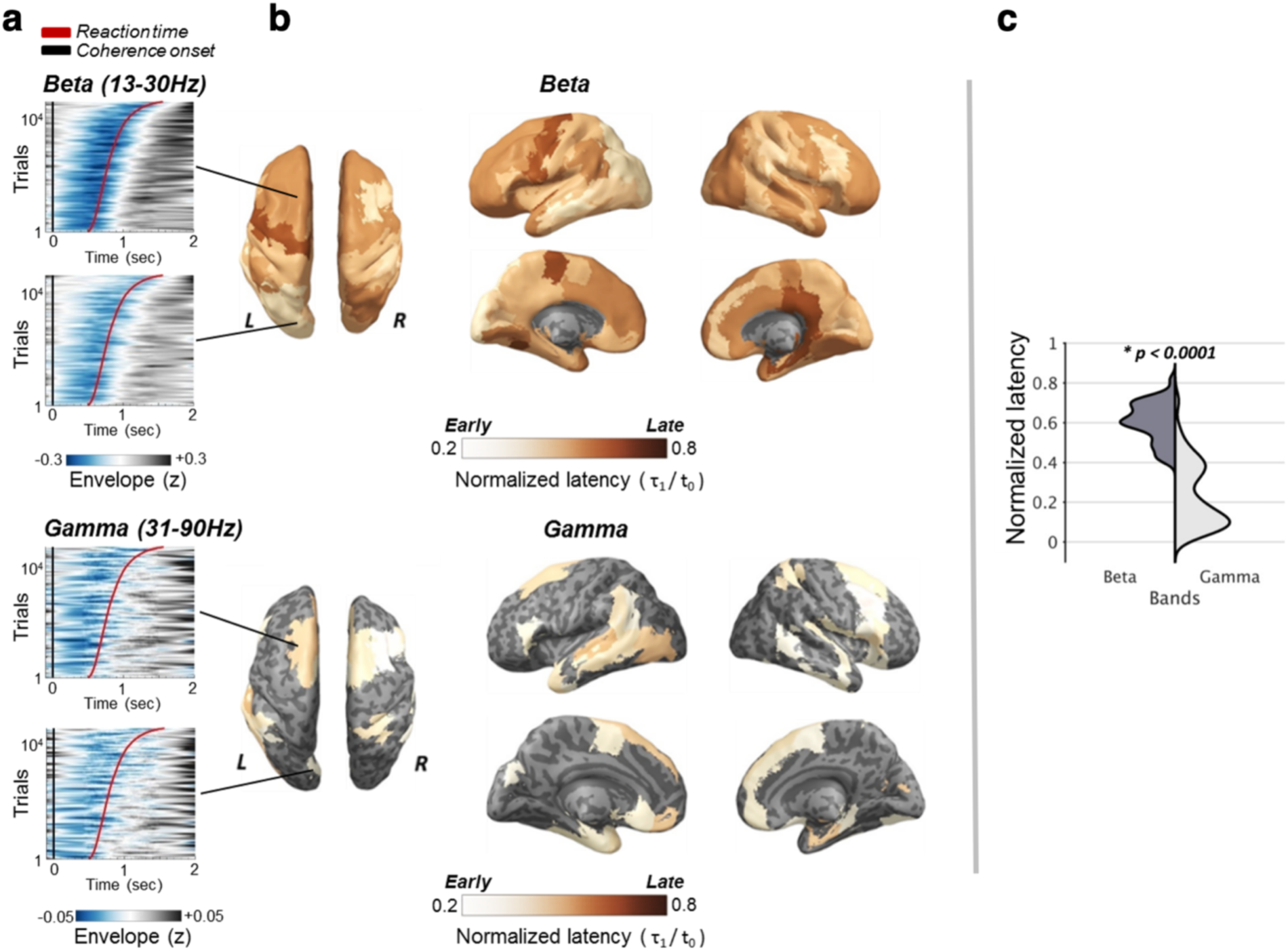
Temporal cascade of decision-evidence accumulation revealed by comparing trial-by-trial MEEG power envelopes to model’s predictions. **(a)** Power plots ranked by reaction-times showing the temporal relationship between signal power envelope (z-scored with regard to each individual’s baseline average) and reaction times (red curve) at representative ROIs showed separately for Beta band (top row) and gamma band (bottom row). The ordinate of each plot represents individual trials pooled across participants and sorted according to reaction times. The black line indicates motion coherence onset. **(b)** Latency maps showing the normalized latencies (each accumulation onset time divided by individual non-decision time) of decision-evidence accumulation across anatomical regions where correlations between power-envelopes and model’s predictions survived random permutation testing (grey areas denote not significant correlation). **(c)** Decision-evidence accumulation in the gamma band precedes beta (top panel).

In the beta band, desynchronization was strongly modulated by uncertainty in good agreement with our predictions. As the decision unfolds, the accumulated decision-evidence will ramp quickly with low perceptual uncertainty, and slowly with high perceptual uncertainty. Accordingly, desynchronization of beta power-envelopes averaged across trials and ROIs was larger (p < 0.0001, cluster corrected random permutations) for low than high perceptual uncertainty^18,29^.

When a response is chosen between multiple options, the race underlying the selection of each alternative is characterized by an overall larger amount of decision-evidence summed across all he racing accumulators by the time of response^13,14^. Accordingly, desynchronization of beta power-envelopes averaged across trials and ROIs was larger for high than low action uncertainty (p < 0.0001, cluster corrected random permutations). Gamma power-envelopes, showed a similar trend, but the effects were statistically insignificant.

Decision-related dynamics expressed by beta desynchronization were distributed across a wide network (Fig3b, mean across significant ROIs: sign-test z = −3.15 ± 0.48, p = 0.00065 ± 0.0016, FDR corrected) similar to previous EEG reports^31^.

In the gamma band we observed a more localized mosaic of ROIs including contralateral motion sensitive areas (inferior lateral occipital region), bilateral extrastriate areas and bilateral frontal motor regions (comprising premotor areas and supplementary motor area; mean across significant ROIs: sign-test z = −2.27 ± 0.27, p = 0.0058 ± 0.003, FDR corrected).

Comparisons between z-transformed correlation values from each of the four levels of our manipulations in isolation confirmed that the quality of fit and results did not vary across trials types (p>0.05, FDR corrected).

### A continuous flow of information

We traced the spectrally resolved temporal evolution of decisions through the visuo-motor hierarchy, finding that decision-evidence accumulation emerges with distinct spatio-temporal profiles between beta and gamma (**Fig. 4b**). An early wave of accumulation begins at ∼120ms from coherence onset within the sparse network oscillating at gamma frequency. It is followed by a second wave mediated by Beta at ∼160ms from coherence onset (**Fig. 4c**; Conjunction of significant ROIs in beta and gamma, median latency across participants, z = 5.53, p<0.0001, Wilkoxon rank test). No difference in latencies was found between hemispheres across frequency bands.

The latency maps (**Fig. 4b**) show an accumulation gradient towards the precentral gyrus. We fitted a piecewise regression model with a free internal knot to the mean latencies of ROIs located along the dorsal path (**Fig. 5a**), a critical system for visuomotor decisions^32,33^. In keeping with our observations the model (**Fig. 5b** left top-bottom panels) identified the precentral gyrus (comprising primary motor cortex and part of the premotor cortex) as the point of convergence of two linear functions (R^2^ = 0.734, p < 0.0001) and outperformed a single regression model (piecewise R^2^ _adj_ = 0.681; linear R^2^ _adj_ = 0.649; adjusted R^2^ penalizes extra free parameters in favor of simple models).

**Figure 5.**
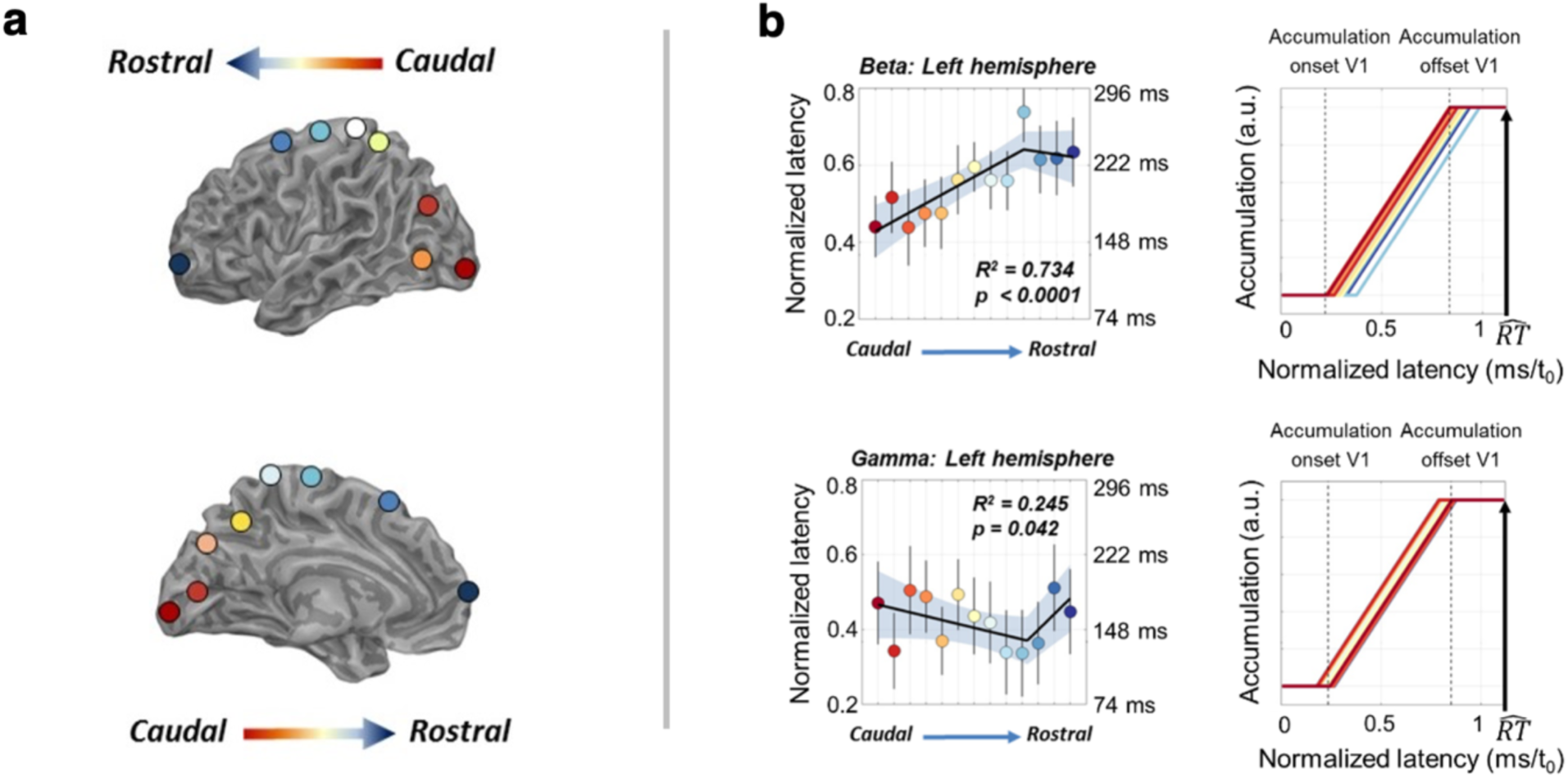
A quasi-parallel cascade of decision-evidence accumulation unfolds along the caudo-rostral axis of the dorsal path. **(a)** Regions of interest (ROIs) along the dorsal path color-coded with respect to their position along the caudo-rostral axis **(b)** Decision-evidence accumulation mediated by beta follows a caudo-rostral gradient along the dorsal path of the contralateral hemisphere. A piecewise regression (top left panel) best describes the gradient showing that latencies increase from visual areas up to M1 in the precentral gyrus and decrease afterwards suggesting two separate converging flows (Error bars indicate SEM, shaded area covers bootstrapped 95% regression CI). The pattern is inverted for gamma where latencies increase while proceeding from M1 to posterior and anterior ROIs. Despite the differences in latencies along the gradient, the cascade of decision-evidence accumulation is quasi-parallel. The right panels show that the latest ROI in the gradient starts accumulating decision-evidence before the earliest ROI (e.g. V1 for beta) has reached the decision boundary. RT hat is the mean reaction time (across trials and participants) normalized by the mean non-decision time (*t*_*0*_).

Interestingly, in the gamma band (**Fig. 5b** bottom left panel) we found a mirror-symmetric trend with increasing accumulation latencies while proceeding from the precentral gyrus to more posterior and anterior regions (R^2^ = 0.245, p = 0.042). Thus, accumulation starts with gamma at ∼120ms from coherence onset in the precentral gyrus and at ∼160ms in the occipital and frontal poles.

The onset of the accumulation in beta overlaps with gamma in the occipital pole at ∼160ms from coherence onset^34^. The interval from earliest onset of accumulation to last onset, is only ∼100ms and the onset in precentral gyrus is on average ∼570ms before a motor response^35^. The delay from motion onset to the beginning of the accumulation on the occipital pole (∼160ms), and the delay from action decision to movement initiation in precentral gyrus (∼100ms) are close to the sensory (∼200ms) and motor (∼80ms) delays measured from neural recordings on macaque^11,36^.

These patterns, albeit with lower spatial resolution, were also found at the sensor level (**SI Appendix, Fig. S4**). As a note of caution for the piecewise regression, the fit of the LBA model for some of the ROIs within the dorsal path was not significant in the gamma band, reducing the accuracy of their latency estimates.

An important observation is that the latest ROIs in the gradient for both beta and gamma starts accumulating decision-evidence before the earliest ROI (e.g. the occipital lobe for beta) has reached its decision boundary (**Fig. 5b** right top-bottom panels). This suggests that decisions are made on the basis of a continuous flow of information, rather than a serial sequence of discrete decisions.

### Information flow from perception to action

The above analyses identified a flow of information across a widespread visuomotor network. To functionally segregate accumulators sub-serving perceptual and action decisions, and to reveal the influx and efflux of information across them we measured the phase-transfer entropy, a data-driven measure of information flow that is robust to signal leakage^22^.

**Fig. 6a** shows, for the beta band, the regions modulated by action uncertainty (Action decision regions, p_corrected_ < 0.0005 in all ROIs) and perceptual uncertainty (Perceptual decision regions, p_corrected_ < 0.0005 in all ROIs). Action decision regions include ipsilateral cingulate and paracingulate cortex^37^, contralateral frontopolar cortex, ventromedial cortex, insula, supplementary motor cortex, inferior parietal lobule and medial parietal cortex^13,38^. Of notice, bilateral precentral gyri were identified as action decision regions which replicates previous findings^13,30^.

**Figure 6.**
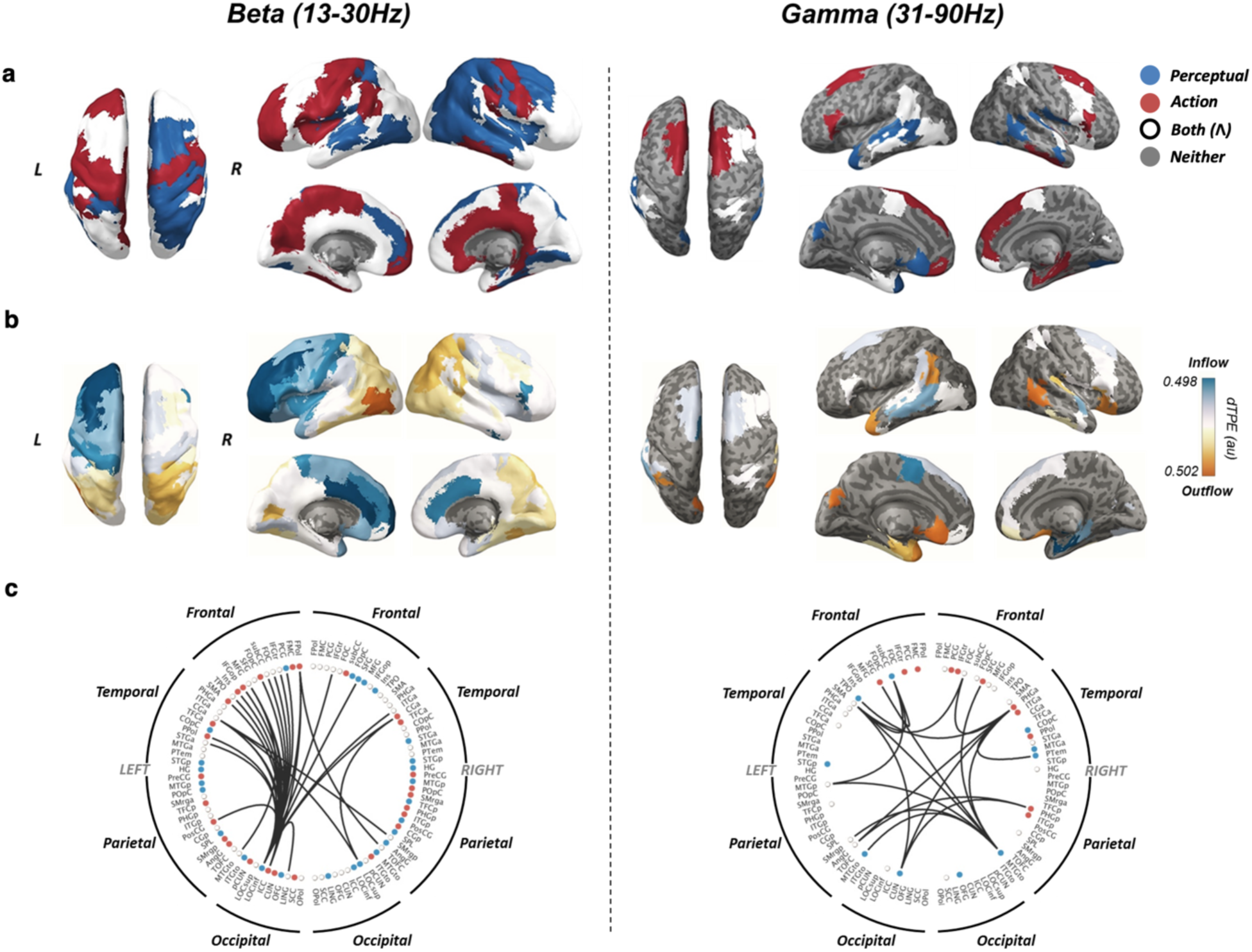
Sensitivity to uncertainty and information flow, show distinct spatiotemporal gradients. **(a)** Differences in phase transfer entropy between manipulations of perceptual and action uncertainty for beta (left column) and gamma (right column) allows to define regions accumulating information specific to perception and action decision. **(b)** Information flow (directional phase transfer entropy) shows a clear rostro-caudal gradient in beta with MT-complex and the frontal regions being the strongest sender and receiver of information, respectively. **(c)** Thresholded connectivity plots. Beta activity reflects transmission of information across distant cortical regions mostly within the left hemisphere. Gamma shows a more local activity with no clear lateralization. The full names of the ROIs are given in **Table S2**.

Perceptual decision regions in the contralateral hemisphere include posterior areas typically associated with decisions about motion direction. These include lateral occipital cortex (including motion area MT-complex), superior temporal cortex (comprising the superior temporal sulcus) and the superior parietal lobule comprising the superior intraparietal sulcus^39^ along with the dorsomedial frontal cortex^4,40^. Interestingly, two areas along the dorsal path on the left hemisphere were sensitive to both perceptual and action uncertainty manipulations (superior frontal gyrus, middle frontal gyrus, lateral occipital cortex superior division (comprising V2 and V3; p_corrected_ < 0.0005 in all ROIs).

In the gamma band, we observed bilateral involvement of the superior frontal gyrus^41^ and inferior frontal gyrus pars triangularis^5^, along with contralateral frontal medial cortex^13^ and ipsilateral paracingulate gyrus in action decisions (p_corrected_ < 0.005 in all ROIs). Perceptual decision areas (p_corrected_ < 0.0005 in all ROIs) included bilateral superior temporal areas (comprising the superior temporal sulcus^39^), cuneal cortex, and subcallosal cortex which has been linked to early encoding of confidence for perceptual decisions^42^.

The dominant direction of information transfer between ROIs was estimated using the directed phase-transfer entropy^22^. The average direction of information flow for each ROI was computed resulting in a single estimate of preferred direction of information flow (either inflow or outflow). Based on these estimates, we calculated a posterior-anterior index^22^ (PAx) to quantify the direction of flow between caudal and rostral ROIs.

**Fig. 6b** show the smooth global pattern of preferential information flow in the beta range with caudal ROIs preferentially sending information to anterior regions. This pattern is similar to that reported previous work in human resting state^22^, except that our results show a task-related lateralization, with the contralateral PAx almost twice the size of the ipsilateral one (left: p = 0.0001, PAx = 0.357; right: p = 0.0083; PAx = 0.184; left vs right PAx, p<0.0001).

It can be seen from **Fig. 6b-c** that, for beta, the strongest information flow was from the left lateral occipital cortex to the left middle frontal gyrus and the frontopolar cortex. This accords with previous reports of beta-synchronization between primate MT and frontal regions during motion discrimination^43^. No significant preferential flow from posterior to anterior regions was seen for the gamma range in either hemisphere (left p = 0.79; PAx = 0.05; right p = 0.37; PAx = 0.3) which is expected given the pattern of accumulation latencies observed in gamma but might also reflect the shorter range of gamma interactions^44^.

### Integration of behavioral, computational and physiological evidence

To highlight the behavioral relevance of the integrated account of visuomotor decision-making, we explored the relationships between connectivity, accumulator model parameters and behavior. To account for multiple-comparisons, we used Holm-Bonferroni correction over eight tests.

In the beta range, the caudo-rostral gradient of evidence-accumulation is matched by a gradual transition from perception to action decisions, as shown by a positive correlation between regional specificity to the type of uncertainty and the estimated accumulation latencies (**Fig. 7a** top left panel, r = 0.27, p_corrected_ = 0.044, 95%CI [0.079 0.420]). Moreover, the information flow is aligned with the caudo-rostral gradient of accumulation since the flow proceeds from perceptual-decision regions to action-decision regions (**Fig. 7a** bottom left panel, correlation between regional specificity and direction of information flow: r = −0.37, p_corrected_ = 0.0016, 95%CI [−0.555 −0.203]).

**Figure 7.**
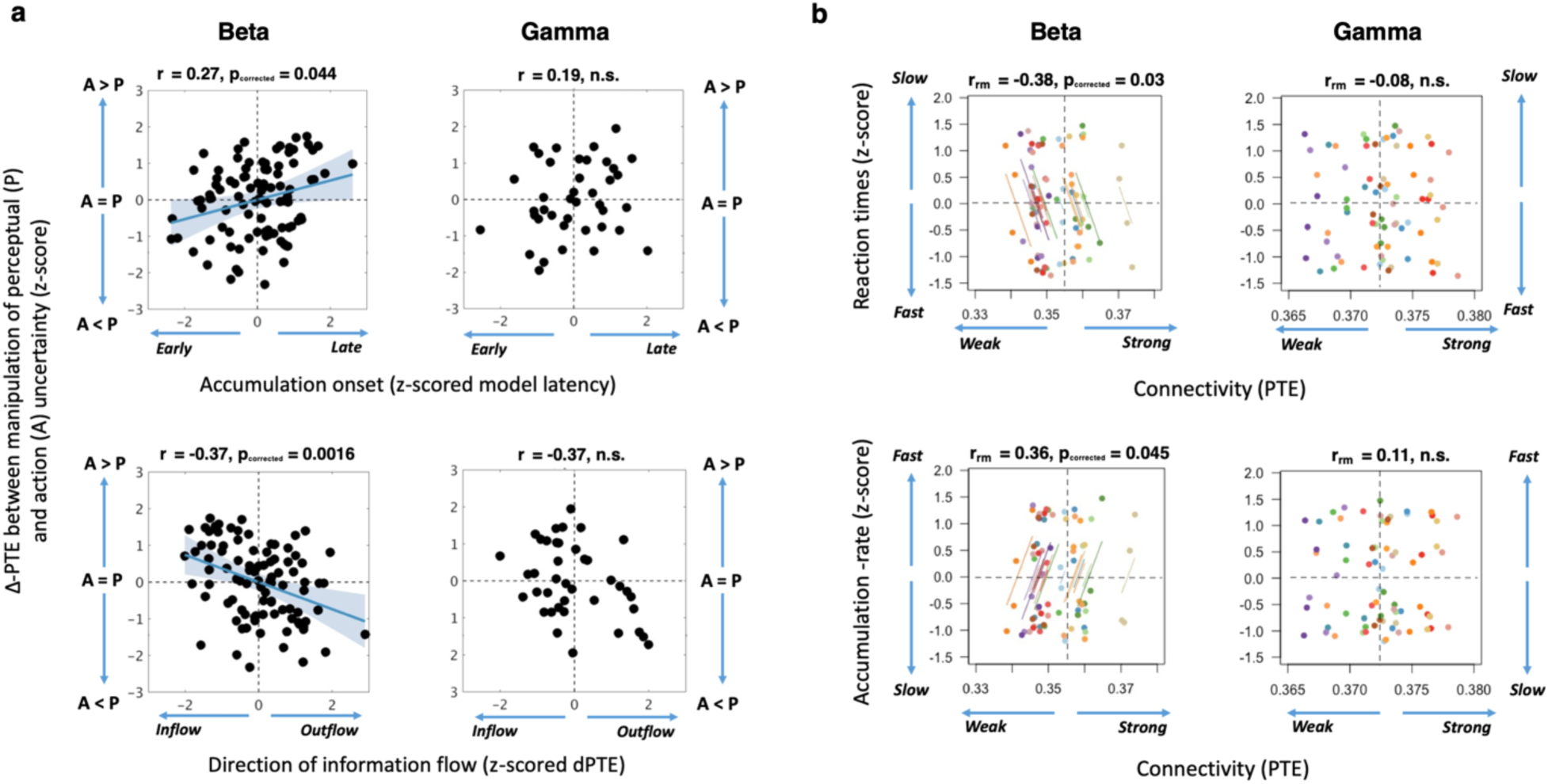
Information flow is stronger under low uncertainty and associated with faster reaction times and larger model accumulation-rates. **(a)** Specificity to uncertainty manipulations correlated with model latency (top row) and direction of information flow (bottom row). Perceptual regions tend to show shorter latencies and to drive information transmission more than action regions. Points show z-scores of median values across subjects for each ROI (96 beta ROIs, 40 gamma ROIs). Shaded area covers bootstrapped 95% CI. **(b)** Repeated-measures correlations. Reaction times (top-row) and accumulation-rates (bottom row) correlated with phase transfer entropy in the beta (left column) and gamma (right column) band. Points show median values for the four tested conditions. Data from the same participant are displayed in the same color, with the corresponding lines showing the individual fit of the repeated-measures correlation

In contrast, for the gamma band we found there was neither significant relationship between region specificity and accumulation latency (**Fig. 7a** top right panel, r = 0.19, p_corrected_ = 0.759) nor significant evidence of flow of information from perception to action decision regions (**Fig. 7a** bottom right panel, r = −0.37, p_corrected_ = 0.068).

Finally, we hypothesized that in a continuous flow of information the amount of information transferred between perceptual and action accumulators co-varies with the rate of accumulation. Faster progression from perception through action should be correlated with phase transfer entropy and model accumulation rate, but negatively with reaction-times.

This was the case in the beta range where strong flow was associated with short reaction times (**Fig. 7b** top left panel, repeated-measures correlation: r_rm_= −0.378, p_corrected_ = 0.03, 95%CI [−0.588, −0.12]) and accumulation rates (**Fig. 7b** bottom left panel, r_rm_= 0.356, p_corrected_ = 0.045, 95%CI [0.096, 0.572]). No significant correlation was observed in gamma (**Fig. 7b** top-bottom right, reaction times vs information flow: r_rm_= −0.079 p_corrected_ = 0.564; accumulation-rate vs information flow: r_rm_ = 0.113, p_corrected_ = 0.816).

## Discussion

There are two principal results from this study that illuminate the interaction between neural systems for perception and action. First, decisions in regions sensitive to motor precision do not wait until regional sensory decisions are completed. Instead, the accumulation of evidence in motor decisions begins within 100ms after the initiation of evidence accumulation in the early sensory regions. This indicates a continuous flow or cascade of information and its gradual transformation from sensory evidence to motor intention^45^. Second, the correlates of evidence-accumulation in the beta and gamma frequency ranges have distinct spatiotemporal profiles, and opposite dominant directions of flow. This spectral directionality is predicted by hierarchical cortical networks for prediction and inference in visuomotor control^18,46–48^. In the beta band, there is not only a spatial gradient in the timing of accumulation-to-threshold between occipital and pre-central cortex, but also a qualitative change in the accumulated signals: from sensitivity to visual uncertainty to sensitivity to response uncertainty. Variability in the strength of the beta-mediated flow of information connecting visual to motor areas, accounts for participants’ decision speed and corresponds to variability in the evidence accumulation rate (**Fig. 7b**). This relationship points to the strength of information flow as a neurophysiological correlate of the speed of evidence accumulation towards a decision.

We set out to integrate the analysis of information flow with decision-making implemented by the accumulation of evidence, and assess their joint influence on trial-to-trial variation in behavior (see **Fig. 1**). Independent manipulation of perceptual and action uncertainty was coupled with the decomposition of performance into latent variables in a parsimonious linear ballistic accumulator model^23^, which accurately generated the response distributions in each task condition including the expected effects of task variance on response latencies^11,14^. The model predictions of within-trial accumulation were correlated with change in beta and gamma power after the onset of stimulus coherence. Beta desynchronization has been shown to scale with uncertainty^30^, but here we show its interaction with the temporal evolution of decision making over sub-second intervals. The observed desynchronization displays two signatures of the accumulation-to-threshold class of models: accumulation of decision-evidence over time and the consistent bound reached shortly before each movement^10,29,35^.

Beta and gamma desynchronization are correlated with behavioural performance. For example, in direct recording from non-human primates during working memory^49^ and sensory discrimination^50^, the beta band desynchronization was greater for accurate trials compared with inaccurate trials. Such beta power encoding of decision outcomes is supramodal in many cortical areas^20^. The change in beta power followed the change in gamma power as in the current study: we found an early wave of gamma followed by a second wave of beta.

Although gamma and beta rhythms have been observed to occur together or in close succession^51,52^, the temporal relationship is functionally relevant. For hierarchical cortical networks, message passing between regions is a function of the laminar asymmetry of afferent vs. efferent connections^46^, and the properties of columnar circuitry which preferentially generates gamma rhythms superficially, and lower frequencies from deep layers^53,54^. This promotes predictive feedback connectivity in beta and lower frequencies, and preferential feedforward ‘error’ signalling in gamma^18,48^. The beta band’s lower frequency makes it inherently more suitable for coordination of information processing over longer conduction delays^33^. As seen in **Fig. 4**, where changes in spectral power were predicted by the LBA model, the latency to accumulation was confirmed as shorter for gamma than beta. Indeed, the spatial distribution of beta latencies in the dominant hemisphere (**Fig. 5b**) shows a gradient from occipital, to parietal and prefrontal, and lastly motor cortex. The motor cortex is also a region of strong net influx of beta (**Fig. 6b**), even more than premotor cortex, consistent with the active inference model of motor control^18,48^.

The spatial gradient of gamma latencies is reversed, with earliest changes observed in precentral cortex, before occipital cortex, and later gamma latencies in time with beta responses in occipital cortex. This may be because of the difference between predicting when a response may be required and what that response should be^55^. The sensory stimulus change (visual coherence) in our task is not the result of the participant’s own response, but is predictable a second after the onset of the non-coherent display. The participant can predict when an action is required, but not which actions are permitted or specified. An increase in localized and predominantly short-range interactions in gamma range may therefore be a permissive of information required for the beta-mediated decision between action alternatives^56^.

Despite the similar latency of beta and gamma accumulators in occipital cortex, the connectivity analyses indicate distinct routing information at longer and shorter spatial scales, respectively. The pattern of net efflux *vs*. influx of beta (**Fig. 6b**) shows a clear division between frontal cortex and posterior lobes. Moreover, the more sensitive a region is to action uncertainty (*vs.* perceptual uncertainty), the later its onset of beta accumulation, and the greater its bias to inflow (*vs.* outflow, **Fig. 7a**). In other words, there was a cascade of overlapping accumulators and information flow along a rostro-caudal axis from perceptual to motor regions for beta, at least in the hemisphere contralateral to the response hand.

Lateralized beta activity during a decision-making task reflects not just movement preparation, but has also been related to a dynamic decision process with updating of a motor plan as a decision evolves^29,30,35^. The beta power lateralization in motor areas was correlated with the state of decision-evidence. Crucially, these earlier MEG and EEG studies used a fixed-mapping between decisions outcomes and categorical behavioural responses, without choice or independence of perception and action decisions. When this fixed mapping between perceptual decisions outcome and motor responses is removed, sensorimotor beta lateralization disappears^57^. Our findings complement this work by directly revealing a lateralized progression of evidence accumulation from posterior perceptual regions to anterior motor areas.

Previous work on visuomotor decisions has often focused on processes occurring at the final choice stage, leaving unresolved the question of whether evidence accumulation is coordinated throughout the cortex or in specific regions. Our findings support a generalized model in which accumulation-to-threshold provides a canonical mechanism evolving throughout all layers of a visuomotor transformation (**Fig. 3a**) and suggest that evidence accumulation is not a limited (perceptual) process with a single cortical focus, but is distributed^58,59^ and applicable to non-sensory evidence or intentions. This multi-focal property of evidence accumulation resonates with results from animal optogenetic^58^ and pharmacological^59^ studies showing that inactivation of local cortical areas carrying decision-related activity did not affect decision-making performance.

Taken together, our observations suggest that the beta band response links sensory evidence to motor plans, throughout a widespread network^60^. We propose that an early neural signalling regarding the need for a response is followed by a second phase that integrates a continuous flow of information to make a decision between them^61^. In this second phase, decisions unfold on the basis of a continuous flow of information (**Fig. 5b**), rather than sequential completion of intermediate decisions at the population level. However, this hypothesis refers to the population level, and we cannot exclude the possibility that within each region, a subsection of neurons completes the relevant decision and forwards this outcome to the next level in the hierarchy, while others in that region continue to accumulate.

The fluctuations in the strength of information flow caused by changes in uncertainty are behaviourally relevant, in their positive correlation with accumulation-rate and negative correlation with reaction times. This establishes an important formal link between behaviour, models of decision-making, and physiological connectivity: Fast accumulating-rates of the linear ballistic accumulator model are associated with a more effective beta-mediated information flow throughout the visuo-motor processing hierarchy, resulting in faster decisions and responses (**Fig.7**). This link provides a powerful tool to investigate clinical conditions in which the ability to use sensory inputs to guide actions is impaired. Examples include Progressive Supranuclear Palsy, Parkinson’s disease and functional movement disorders, which are associated with abnormal evidence accumulation rates^62,63^, and abnormal beta power^64,65^.

In summary, our analytical approach set out in **Fig 1** elucidates behavioural decisions through the combination of computational modelling of behaviour and identification of neurophysiological signatures of latent variables, in distributed cortical networks. Beta power reflects the temporal and spatial dynamics of the accumulation and transfer of decision-evidence, with a continuous flow of information between regions rather than sequential discrete decisions. During this flow, the gradual transition from the resolution of sensory uncertainty to the resolution of response uncertainty, enables goal-directed actions in the face of uncertainty.

## Materials and Methods

### Participants

Twenty healthy volunteers (9 females, 11 males, age range 18-39 years) took part in this study, after providing informed consent. Inclusion criteria included age 18-40 years, right-handed, and screening for neurological or psychiatric illness. Two subjects failed to reach the requisite performance criterion during training and were excluded, leaving 18 subjects in all subsequent behavioral and neural analyses. The chosen sample size was based on previous EEG/MEG studies on perceptual and motor decisionmaking^35,66^. Experimental protocols conformed to the guidelines of the Declarations of Helsinki and were approved by the local research ethics committee.

### Task and procedures

Participants performed a finger-tapping task adapted from previous studies ^13,14^. Their goal was to detect the onset of coherent motion and to press the button corresponding to one of the downward moving stimuli (coherent stimuli). The number of coherent stimuli defined two trial types: Low action uncertainty trials, where a single coherent stimulus commanded which button to press; and high action uncertainty trials, where three coherent stimuli required the participants to make a simple choice and press any one of the three corresponding buttons (a “fresh choice, regardless of what you have done in previous trials”^14^). Equal emphasis was placed on the speed and accuracy of the responses. Participants were instructed to fixate on a central red mark throughout the trial. Eye-tracking data collected during the first six scanning sessions confirmed participants were able to successfully perform the task while maintaining fixation (see **SI Appendix, Fig. S5**). Each trial started with the presentation of the fixation mark and stimuli onset ensued after a variable interval comprised between 0.5sec and 1sec. The imaging session was preceded by one training psychophysical session and one test session scheduled on separate days; the scanning session was conducted a maximum of four days after the psychophysical training, depending on the availability of the participants. The test session was to ensure that the participants were able to perform well under the individually adjusted motion thresholds. In the test and scan sessions, coherence levels were fixed to the individual thresholds corresponding to high and low levels of perceptual uncertainty, the match-to-sample task was removed, and no feedback was provided except for too late or too long responses. Levels of perceptual and action uncertainty where randomly interspersed across trials. Each session consisted of 10 blocks (total 720 trials per participant) separated by a short rest. Trials on which responses were made before 0.1-sec or after 2-sec (on average 1.3% of total trials) were excluded from subsequent analyses.

### Stimuli

Stimuli were presented using Matlab (Matworks, Natiek, MA, USA) and the Psychotoolbox routines in a sound-proof and dimly lit room. For the psychophysical training stimuli were displayed on a CRT monitor at 60cm, and for the scan session stimuli were projected on a screen through a projector at 130cm (both with a 60Hz refresh rate) with equivalent pixel resolution of 0.03°.

Stimuli were four random dot kinematograms displayed within four circular apertures (4° diameter) positioned along a notional semi-circular arc (3.4° eccentricity) on a black background (100% contrast). 200 dots were displayed during each frame and spatially displaced in the next frame to introduce apparent downward motion (6°/sec velocity). To manipulate motion strength (i.e. motion coherence) between trials, on each frame only a certain proportion of dots moved downward whilst the rest of the dots where randomly reallocated. Motion coherence level was kept constant throughout the trial.

Since abrupt stimulus onset and offset could elicit large sensory-evoked potentials which might mask decision processes, the 1.5 seconds long coherent motion interval was preceded and followed by intervals of zero-coherence levels lasting 1sec and 0.5sec, respectively.

### Psychometric calibration

Participants were firstly familiarized with the finger-tapping task during a short practice session where 100% coherent stimuli were adopted. The familiarization phase was completed when participants reached 90% accuracy across all trial types. In the following psychophysical training, motion coherence was randomly varied between trials to estimate individual motion thresholds. Eight logarithmically spaced motion coherence levels (0 0.5 0.10….0.9) were used (32 trials per level) following extensive piloting to ensure coverage of a wide range of individual motion sensitivity. Each training session comprised 16 blocks of 32 trials. Feedback was provided for correctness of responses as well as for too early or too late responses (100ms and 2.5s from motion coherence onset, respectively).

To ensure that participants perceived all the available options (i.e. coherent stimuli) before committing to a decision, occasionally (p = 0.2) after a correct choice they had to perform a secondary match-to-sample task: a set of grey discs replaced the stimuli and participants had to report whether their locations matched the location of the previously displayed coherent stimuli. They had to press any button to report a match and withhold any response otherwise. A trial was considered as correct only when both choice and matching were correct. Trials with un-matching responses were discarded and repeated within the session.

To tailor the sensory evidence to the participants’ individual motion sensitivity across number of options, the discrimination accuracy of each trial type in each training session was fitted using a maximum likelihood method, with a Log-Quick function defined as

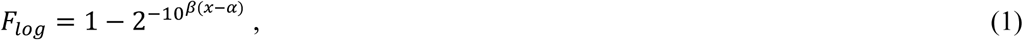

where α is the threshold, β is the slope and x is the coherence level. To obtain the proportion correct for each trial type, the Log-Quick function was scaled by,

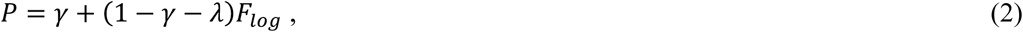

where γ is the guess rate and λ is the lapse rate controlling the lower and upper asymptote of the psychometric function, respectively.

Individual low and high perceptual uncertainty levels for each trial type were estimated as the 75^th^ and 90^th^ percentile of the psychometric functions from the last session. The reason for adopting these thresholds was twofold: firstly, participants need to perceive all the available options before committing to a decision. Secondly, supra-threshold trials are best suited for investigating neural correlates of evidence accumulation ^67^.

### MEG and EEG data acquisition and processing

An Elekta Neuromag Vectorview System (Helsinki, Finland) simultaneously acquired magnetic fields from 102 magnetometers and 204 paired planar gradiometers, and electrical potential from 70 EEG electrodes (70 Ag-AgCl scalp electrodes in an Easycap - GmbH, Herrsching, Germany - extended 10-10% system). Additional electrodes provided a nasal reference, a forehead ground, paired horizontal and vertical electro-oculography, electrocardiography and neck electromyography. All data were recorded and digitized continuously at a sample rate of 1kHz and high-pass filtered above 0.01 Hz.

Before scanning, head shape, the locations of five evenly distributed head position indicator coils, EEG electrodes location, and the position of three anatomical fiducial points (nasion and left and right pre-auricular) were recorded using a 3D digitizer (Fastrak Polhemus Inc., Colchester, VA). The initial impedence of all EEG electrodes was optimized to below 10 kΩ, and if this could not be achieved in a particular channel, or if it appeared noisy to visual inspection, it was excluded from further analysis. The 3D position of the head position indicators relative to the MEG sensors was monitored throughout the scan.

Environmental noise suppression, motion compensation, and Signal Source Separation for the MEG data was followed by separate independent component analysis for the three sensor types and artifactual components were rejected. For EEG data, components temporally and spatially correlated to eye movements, blinks and cardiac activity were automatically identified with EEGLab’s toolbox ADJUST (Swartz Center for Computational Neuroscience, University of California San Diego). For MEG data, components were automatically identified that were both significantly temporally correlated with electrooculography and electrocardiography data, and spatially correlated with separately acquired topographies for ocular and cardiac artifacts using in-house Matlab code.

The continuous artefact-corrected data were low-pass filtered (cut-off = 100Hz, Butterworth, fourth order), notch filtered between 48 and 52Hz to remove main power supply artifacts, down-sampled to 250Hz, and epoched from −1500 to 2500ms relative to motion coherence onset. EEG data were referenced to the average over electrodes.

MEG and EEG data were combined before inversion into source space ^21^ using the Miminum Norm algorithm as implemented by SPM12. Notably, combined MEG and EEG allows a better localization of neural sources than each technique on its own^21^. The forward model was estimated from each participant’s anatomical T1-weighted MRI image. All conditions were included in the inversion to ensure an unbiased linear mapping. The source images were spatially smoothed using an 8 mm FWHM Gaussian kernel.

### Accumulator model of perceptual and action decisions

We adopted formal models of decision-making to decompose the behavioral performance into cognitively relevant latent variables. We fitted accumulation-to-threshold models (Linear Ballistic Accumulator^23^, LBA), to each participant’s reaction time and accuracy data. The LBA model of decisions is more tractable than drift-diffusion models for n-way decisions while still remaining physiologically informative^68^ and has been previously applied to a finger tapping task to model fMRI evidence accumulation^13^. Instead of adopting a two-stage model, which assumes a discrete serial process between perceptual and action decisions, we opted for a ‘unitary’ model where both perceptual and action uncertainty concur in determining participant’s performance in a given trial. The factorial design of the experiment enabled us to divorce perceptual and action decision processes using connectivity metrics.

According to this class of models, each accumulator linearly integrates the decision-evidence (or the intention) over time in favor of one action, and the decision is made when the accumulated activity reaches threshold (see Fig.3c). In our task possible actions correspond to a button press from one of four fingers, each modeled by independent accumulators *i* ∈{1, 2, 3, 4}. When three valid actions are available, three accumulators are engaged with activation starting at levels independently drawn from a uniform distribution [0, c0], and increasing linearly over time with an accumulation rate (*v*) drawn from an independent normal distribution with mean μ_i_ and standard deviation σ_i_. A response is triggered once one accumulator wins the ‘race’ and reaches a decision bound (*b).* When only one action is available, only the accumulator corresponding to the available action is engaged. Predicted reaction time (RT) is given by the duration of the accumulation process for the winning accumulator, plus a constant non-decision time *t*_*0*_ representing the latency associated with stimulus encoding and motor response initiation^23^.

To identify the combinations of free parameters that best accounted for the observed behavioral data we firstly fitted 15 variants (i.e. all possible combinations without repetition) of the LBA. Each variant was characterized by a unique combination of *free* parameters allowed to vary across trials. The best-fitting parameters for each model variant were used to calculate the Bayesian Information Criterion (BIC), which penalizes extra free parameters in favor of simpler models. BIC values were then used to compare the goodness-of-fit of each variant using random-effects Bayesian model comparison ^24,25^. Predictions of decision-related activity were generated from the winning LBA model to locate neural signatures of decisions-evidence accumulation in single-trial analyses of MEEG data.

### Estimation of expected neural activity

We generated predictions of decision-related activity from the LBA model^13^ to locate neural signatures of decisions-evidence accumulation in single-trial analyses of MEEG data. For multiple options, the LBA model assumes multiple active accumulators, one for each finger option. Let 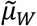 be the accumulation rate of the winning option (i.e. the one reaching response threshold *b*), sampled from the normal distribution 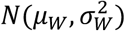. Let 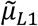 and 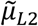 be the sampled accumulation rates of the alternative options (i.e. the losers), sampled from normal distributions 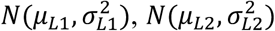, respectively. If the reaction time of a given trial is RT, the latency of the accumulation process is *RT* − *t*_0_, such that the expected accumulation of the winning option is:

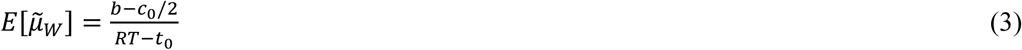

Since the losing accumulators have not reached the threshold by the time of the response RT, the expected values of 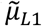 and 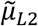 are smaller than 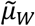.

Therefore, the losing accumulation rates have truncated normal distributions with an upper bound of 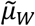 and with expected values of:

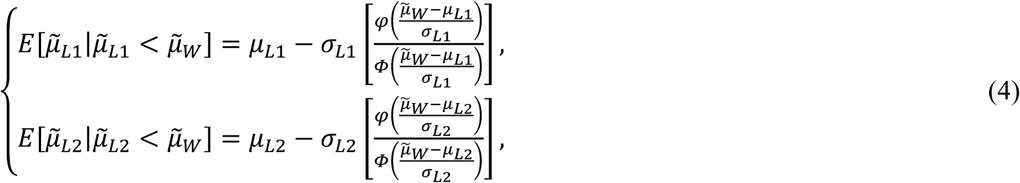

where 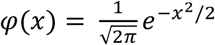 and 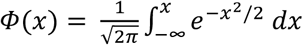.

The sum of the winning and losing accumulation rates gives an estimation of total accumulation activity for single trials. For trials with only one available option, the accumulation activity is determined by the only active accumulator.

### Dimensionality reduction

To improve computational efficiency, reduce multiple comparisons issues while retaining the maximum amount of information^69^, we reduced the dimensionality of the MEEG data by parcellating the cortical surface into a set of 96 regions of interest (ROIs) defined using the Harvard-Oxford cortical atlas (FSL, FMRIB, Oxford) and by representing the dynamic of each ROI with a single time-course, obtained using principal component analysis^70^. The reconstructed sources within each ROI were first bandpass-filtered in either beta (13-30 Hz) or gamma (31-90Hz) frequency bands. The coefficients of the principal component accounting for the majority of the variance of the vertices within each ROI, were then taken as an appropriate representation of source activity for that region. Next, to estimate the power oscillations on a single-trial basis, we extracted frequency-specific signal envelope modulations using a Hilbert transform of the source data from each reconstructed ROI (epochs from 500ms before to 1500ms after coherence onset). The Hilbert’s envelope is a convenient measure of how the power of the signal varies over time in the frequency range of interest, and thus particularly suited to capture relatively slow fluctuations associated to the instantaneous accumulation of evidence/intentions. The power estimates of individual participants were down-sampled to 100Hz and normalized by their baseline (from 400ms to 100ms before coherence onset).

### Single-trial analysis

To identify the spatio-temporal profile of decision-related accumulation over the brain we derived model-predicted signals for each trial to compare with neural oscillations in beta and gamma frequency bands. We estimated the maximum lagged absolute Spearman correlation between the model predicted activity and the signal envelope in a trial-by-trial fashion. The lagged correlation was used to optimally split the non-decision time before and after the accumulation period to determine the time delay between the neural signal and the model predictions. The time before accumulation provides a measure of the temporal separation between coherence onset and accumulation onset.

If the model prediction *x* is a lagged version of the neural signal *y* so that

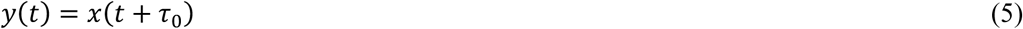

Where *τ*_*0*_ is a time delay that can vary from 0ms to the individual non-decision time (*t*_0_) with steps of 10ms, then the maximum absolute lagged correlation between *x* and *y* is defined as

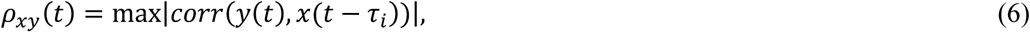

where *i* = [0, 10, 20 … *t*_0_].

With the peak value of ρ_*xy*_(*t*) occurring when τ_*i*_ = τ_0_ which allows us to determine the time delay. We estimated the largest absolute lagged correlation value for each ROI and individuals by comparing concatenated epochs and model predictions. This choice permits to measure accumulation lags specific to each ROI, under the assumption that they differ across brain regions for each participant. The strength of the Fisher-transformed maximum lagged correlations for each ROI was then quantified (z-score) using a one-sample sign-test. To provide a conservative estimate of significant correlations between model prediction and neural activity, we repeated the above procedure 10.000 times, each iteration using a different phase-randomized version of the original MEEG signal, to obtain a distribution of correlations under chance. Two-tailed statistical significance was assessed by computing the proportion of absolute values of the distribution of correlations generated by chance exceeding the correlation between model predictions and the original MEEG signal. The resulting p-values were corrected for multiple comparisons (False Discovery Rate) across ROIs and frequency bands.

### Connectivity analyses

To explore the direction of the information flow we employed phase-transfer entropy, a data-driven effective-connectivity measure robust to signal leakage^22^. The preferred direction of information between ROIs whose activity best matched with model’s predicted activity was estimated using the directed phase-transfer entropy. The analyses focused on regions whose activity significantly fitted the LBA model’s prediction.

To identify the ROIs that preferentially accumulated evidence for perception or action decisions, the average information flow (quantified by phase transfer entropy) sent by each ROI was calculated for each subject and condition. The difference of information flow between uncertainty levels for perception and action is compared at the ROI level with a surrogate distribution generated by flipping the condition labels for a random number of participants (10.000 iterations). Since significance was estimated separately for perception and action, the critical value for the FDR correction was halved to *α* = 0.025.

To quantify the direction of information flow, we calculated a posterior to anterior index (PAx) as implemented by Hillebrand et al, 2016. A positive PAx indicates preferential flow from posterior regions toward anterior regions. ROIs were split into anterior and posterior region with respect to the central sulcus (see **SI Appendix, Table S2**). Significance was assessed with permutation testing where the average directional phase-transfer entropies were shuffled across ROIs and PAx was estimated. This procedure was repeated 10.000 times to generate a surrogate distribution of PAx values against which the observed PAx values were tested (p<0.025 to account for multiple comparisons).

For the correlations in **Fig. 7a**, we first confirmed homoscedasticity of our data and then calculated bootstrapped Pearson’s correlations. For the correlations in **Fig. 7b** we used repeated-measures correlation (as implemented in the rmcorr package in R) which accounts for non-independence among observations due to multiple measurements per participant. The resulting p-values were corrected for multiple comparisons by applying Holm-Bonferroni correction.

### Hypothesis testing

Differences in reaction times were tested with a 2-way repeated measures ANOVA (Low/High Uncertainty x Action/Perception). All other hypothesis tests used non-parametric tests or random permutation methods that do not rely on specific assumptions about the distributions of data values. All tests were evaluated at the p<0.05 level (two-tailed), correcting for multiple comparisons where appropriate.

### Data and code availability

Additional data related to this study are available from the corresponding author upon request. The code and data to generate the results and the figures of this study are available at https://github.com/ale-tom/MEEG_Uncertainty.

## Supporting information

Supplementary results

