## Supplementary results for "On the evolution of neural decisions from uncertain visual input to uncertain actions"

### Supplementary Figures and Tables

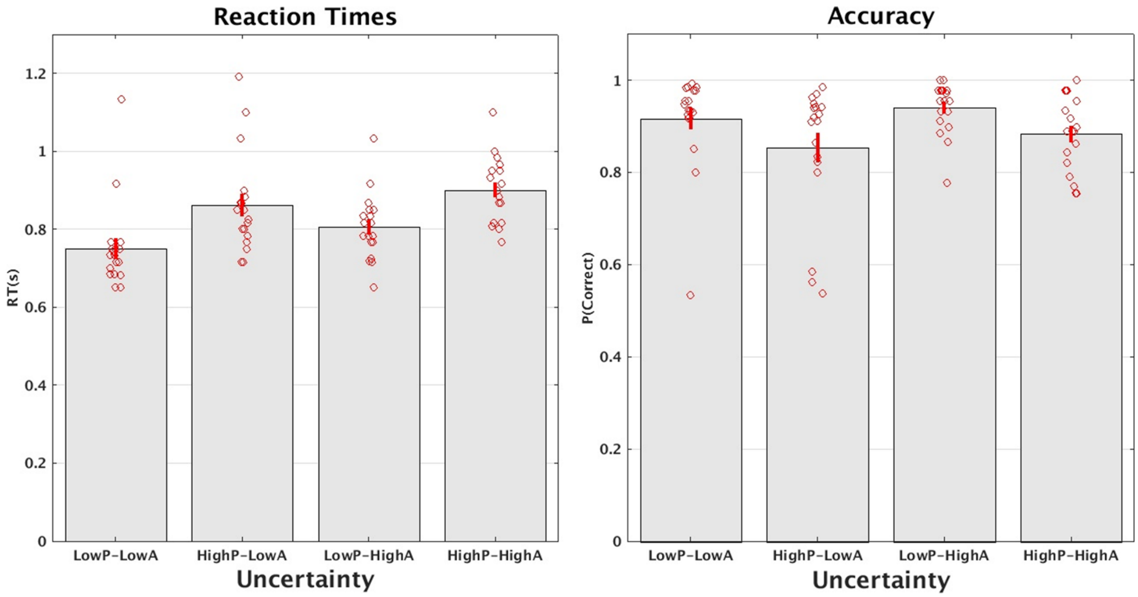


**Figure S1 Behavioural performance during scan**. Reaction times (left panel) for each manipulation level. Accuracy levels (right panel) for each manipulation level. To compare between conditions with different number of options (i.e. low and high action uncertainty) accuracy is normalised with respect to the probability to respond correctly when guessing (i.e. chance level) as follows: (accuracy – chance)/(1-chance). Chance levels are 25 % and 75% for low and high uncertainty, respectively. Red dots indicate individual data, error-bars represent standard error of mean.

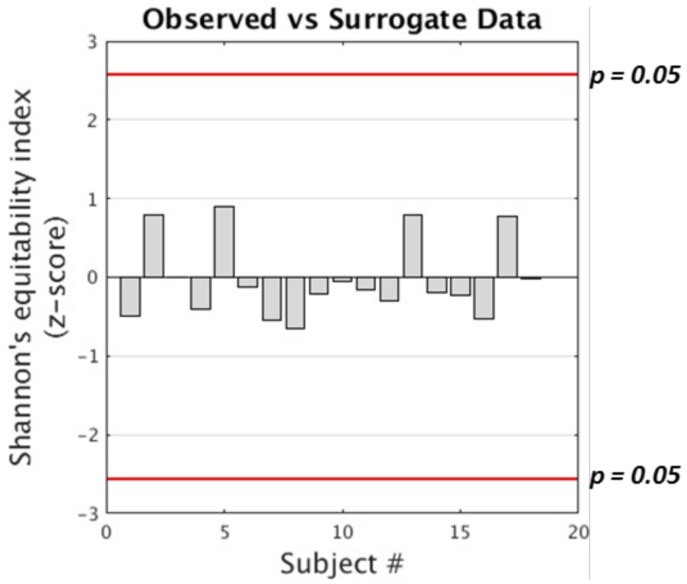


**Figure S2. Shannon's equitability index** quantifies how much a given choice is biased by previous responses. Permutation tests showed that Shannon's equitability index estimated from participants' responses did not differ significantly from that generated by random permutations of trials order **(**red lines show the Bonferroni-corrected significance threshold of p = 0.05; two-tailed). This confirms that over a series of trials subjects' choices were not biased by previous responses.

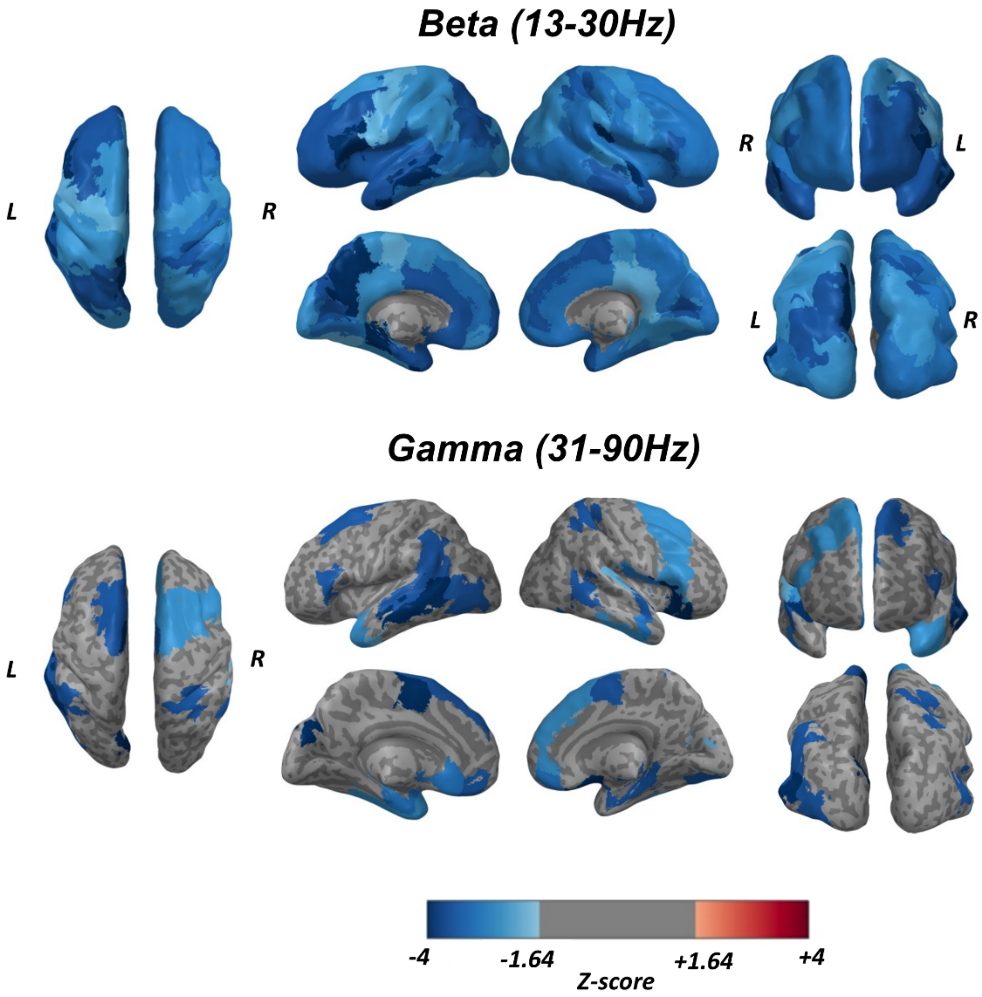


**Figure S3. Correlation between trial-by-trial predictions and MEEG activity in source space.** Colours indicate the overall z-score for difference of empirical correlations from chance in the Beta (top panel) and Gamma (bottom panel) frequency ranges. Empirical correlations were compared against surrogate correlation distributions obtained by correlating predictions to phase randomized MEEG signals (10000 randomization for each ROIs). Grey indicates non significant difference between empirical and surrogate correlations at $p<0.05$ FDR corrected for multiple comparisons.

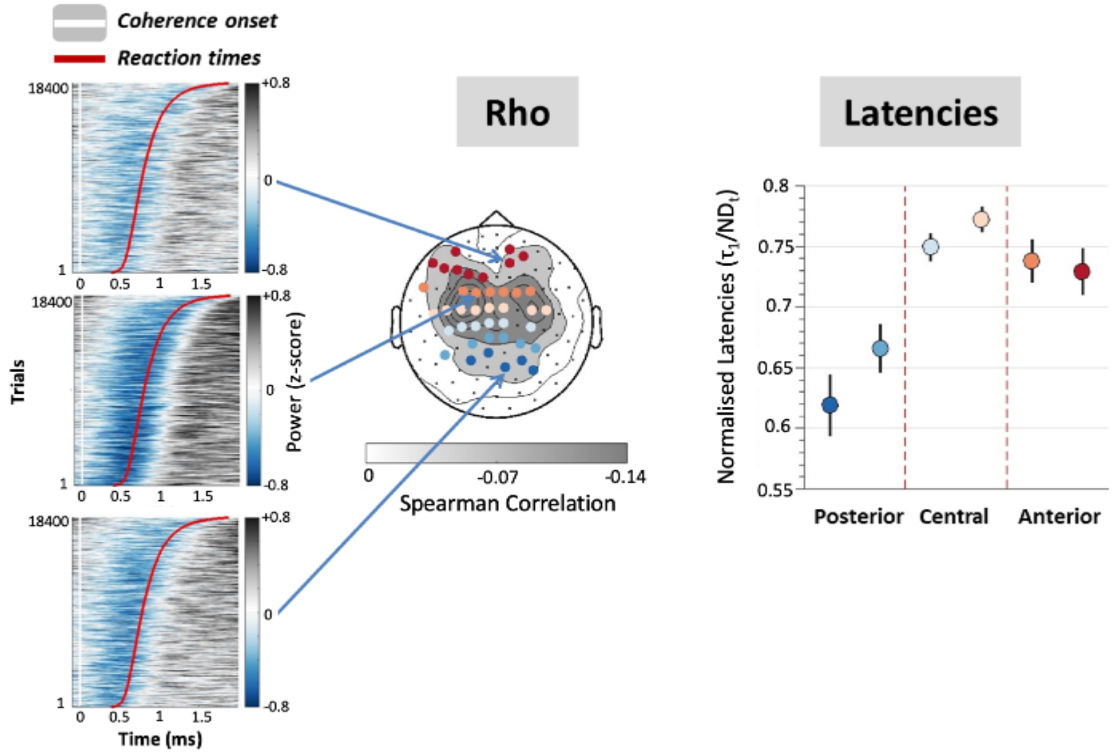


**Figure S4.** Temporal cascade of information-representation in beta (planar gradiometers in sensor space). Similarly to figure 4, the left panel shows power plots ranked by reaction times (red line). The topographic plot in the middle shows sensors where correlations between power-envelopes and model's predictions survived random permutation testing. Sensors are coloured depending on their position along the caudo-rostral axis. The right panel shows a gradient of latencies that increases from posterior areas up to central regions and decreases afterwards. No significant correlations were observed in the gamma band.

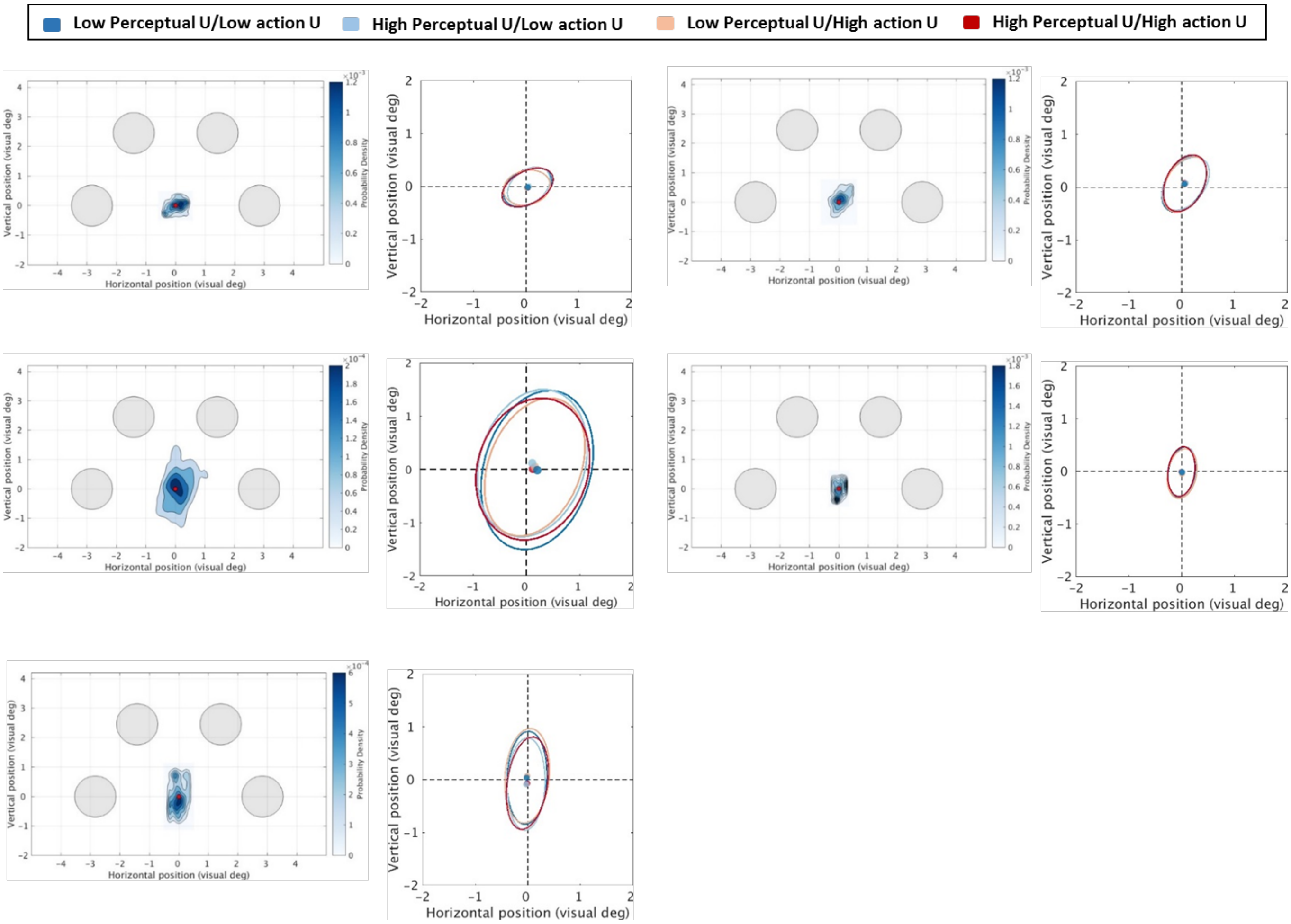


**Figure S5. Eye-tracking data from 5 participants** show that fixation could be easily maintained during the scan session (heat-maps). In the plots on the right hand confidence ellipses encompassing 95% of fixations population are shown for each condition. The colored points indicate the location of the ellipses' centroids. The semimajor and semiminor axes and the orientation of the ellipses were computed from the first and second principal components of the fixations scatter. The position of the ellipses' centroid with respect to the fixation point (centre of cross-hair) gives a measure of fixation accuracy, whereas its area provides an overall measure of fixation dispersion. Neither accuracy nor dispersion of subjects' fixation differed between conditions (2 perceptual uncertainty x 2 action uncertainty repeated-measures ANOVA. Accuracy: perception [$F_{1,4}$(0.379) p = 0.57] action [$F_{1,4}$(0.013) p = 0.91]; Dispersion: perception [$F_{1,4}$(0.078) p = 0.79] action [$F_{1,4}$(0.587) p = 0.48])

| Subj # | Boundary (***b***) | Accumulation Rate (***v***) | | | | Accumulation Rate StDev | Non-decision time ($t_0$) |
|---|---|---|---|---|---|---|---|
| | | *Uncertainty Levels (* ***p*** *:perceptual* ***a*** *:action)* | | | | | |
| | | **LpLa** | **HpLa** | **LpHa** | **HpHa** | | |
| **1** | 1.73 | 7.96 | 5.42 | 4.48 | 2.06 | 4.63 | 0.51 |
| **2** | 4.34 | 18 | 11.7 | 5.6 | 3.47 | 5.59 | 0.42 |
| **3** | 8.03 | 18.33 | 15.54 | 7.08 | 6 | 5.62 | 0.28 |
| **4** | 3.73 | 14.26 | 11.46 | 4.65 | 2.47 | 6.53 | 0.4 |
| **5** | 5.02 | 13.99 | 9.49 | 4.52 | 3.27 | 6.21 | 0.37 |
| **6** | 4.94 | 9.98 | 5.86 | 6.36 | 5.42 | 4.36 | 0.31 |
| **7** | 4.79 | 8.77 | 6.94 | 3.65 | 3.1 | 4.66 | 0.42 |
| **8** | 2.77 | 5.46 | 4.02 | 2.91 | 2.18 | 3.12 | 0.37 |
| **9** | 2.94 | 5.86 | 4.72 | 3 | 2.59 | 4.56 | 0.39 |
| **10** | 5.13 | 11.55 | 9.64 | 6.7 | 5.01 | 4.76 | 0.25 |
| **11** | 1.29 | 4.16 | 3.5 | 2.15 | 1.22 | 3.28 | 0.39 |
| **12** | 2.05 | 5.68 | 3.94 | 2.41 | 1.54 | 3.98 | 0.43 |
| **13** | 4.23 | 10.59 | 8.13 | 4.8 | 3.36 | 4.25 | 0.35 |
| **14** | 6.23 | 6.65 | 5.63 | 5.88 | 5.23 | 6.72 | 0.3 |
| **15** | 2.33 | 6.54 | 5.28 | 2.74 | 1.53 | 4.62 | 0.38 |
| **16** | 1.33 | 5.98 | 4.41 | 1.39 | 1.3 | 3.01 | 0.5 |
| **17** | 3.5 | 14.66 | 11.74 | 3.79 | 2.27 | 5.65 | 0.47 |
| **18** | 8.78 | 17.52 | 14.99 | 8.65 | 8.13 | 4.96 | 0.2 |
| **Mean** | 4.06 | 10.33 | 7.91 | 4.49 | 3.34 | 4.81 | 0.37 |
| **Std Dev** | 2.13 | 4.74 | 3.85 | 1.94 | 1.90 | 1.09 | 0.08 |

**Table S1**. **Parameters for the winning LBA model**

| Acronym | Anatomycal label |
|---|---|
| Opol | **Occipital Pole** |
| SCC | **Supracalcarine Cortex** |
| LING | **Lingual Gyrus** |
| OFG | **Occipital Fusiform Gyrus** |
| CUN | **Cuneal Cortex** |
| ICC | **Intracalcarine Cortex** |
| LOCinf | **Lateral Occipital Cortex, inferior division** |
| LOCsup | **Lateral Occipital Cortex, superior division** |
| pCUN | **Precuneous Cortex** |
| ITGto | **Inferior Temporal Gyrus, temporo-occipital part** |
| MTGto | **Middle Temporal Gyrus, temporo-occipital part** |
| TOFC | **Temporal Occipital Fusiform Cortex** |
| AngG | **Angular Gyrus** |
| SMrgp | **Supramarginal Gyrus, posterior division** |
| SPL | **Superior Parietal Lobule** |
| CGp | **Cingulate Gyrus, posterior division** |
| PosCG | **Postcentral Gyrus** |
| ITGp | **Inferior Temporal Gyrus, posterior division** |
| PHGp | Parahippocampal Gyrus, posterior division |
| TFCp | Temporal Fusiform Cortex, posterior division |
| SMrga | Supramarginal Gyrus, anterior division |
| POpC | Parietal Operculum Cortex |
| MTGp | Middle Temporal Gyrus, posterior division |
| PreCG | Precentral Gyrus |
| HG | Heschls Gyrus (includes H1 and H2) |
| STGp | Superior Temporal Gyrus, posterior division |
| PTem | Planum Temporale |
| MTGa | Middle Temporal Gyrus, anterior division |
| STGa | Superior Temporal Gyrus, anterior division |
| Ppol | Planum Polare |
| COpC | Central Opercular Cortex |
| TFCa | Temporal Fusiform Cortex, anterior division |
| CGa | Cinfulate Gyrus, anterior division |
| ITGa | Inferior Temporal Gyrus, anterior division |
| PHGa | Parahippocampal Gyrus, anterior division |
| SMA | Juxtapositional Lobule Cortex (formerly Supplementary Motor Area) |
| TPO | Temporal Pole |
| Ins | Insular Cortex |
| IFGop | Inferior Frontal Gyrus, pars opercularis |
| MFG | Middle Frontal Gyrus |
| SFG | Superior Frontal Gyrus |
| FOpC | Frontal Operculum Cortex |
| subCC | Subcallosal Cortex |
| FOC | Frontal Orbital Cortex |
| IFGtr | Inferior Frontal Gyrus, pars triangularis |
| PCG | Paracingulate Gyrus |
| FMC | Frontal Medial Cortex |
| Fpol | Frontal Pole |

**Table 2S**. **Abbreviations for the cortical regions in Figure 5c.** Regions in boldface formed the set of posterior regions (ROI centroids MNI coordinate Y ≤ 42; Y(42) = Postcentral Gyrus) defined with respect to the central sulcus. Thus defined posterior and anterior regions were used to calculate the posterior to anterior index (PAx) to quantify the posterior-to-anterior pattern of information flow.
